# Testing the application of plasma glucocorticoids and their ratios as biomarkers of acute and chronic stress in rescued wild koala patients: a pilot study

**DOI:** 10.1101/2024.05.24.595853

**Authors:** Liang-Yu Pan, Harsh Pahuja, Tim Portas, Edward Narayan

## Abstract

Koalas *(Phascolarctos cinereus)* are one of the most iconic marsupial species endemic to Australia. However, their population is declining due to threats including habitat loss, disease, dog attacks, and vehicle collisions. These threats also serve as acute or chronic stressors that impact koala welfare and conservation. Cortisol is widely used as a biomarker to study stress in koalas. However, plasma cortisol concentration is less studied due to its limited ability to assess chronic stress and welfare concerns. Dehydroepiandrosterone sulphate (DHEAS) and dihydrotestosterone (DHT) are biomarkers that could potentially detect chronic stress due to their antagonising and inhibitory effects on cortisol. In this study, we used plasma cortisol and the ratio of DHEAS and DHT to cortisol to assess stress in rescued koalas (n = 10) admitted to RSPCA Queensland. Although no significant differences were found between koalas across all biomarkers and the ratios failed to detect chronic stressors, similar trends were found consistently, suggesting the potential use of the biomarkers to assess stress. Across all biomarkers, the highest medians were found in koalas with Chlamydia-related reproductive disease and oxalate nephrosis and the lowest medians were found in koalas with Chlamydia-related conjunctivitis. Higher medians were also found consistently in females (n = 3) and adult koalas. In addition, insignificant negative correlations were found across all biomarkers between age, weight, and body conditioning scores, except for the positive correlation between weight and cortisol and cortisol:DHT. Overall, the consistency of trends and the insignificant differences found across biomarkers in our study suggested that using a single biomarker to assess chronic stress is insufficient, especially for hospital-based studies limited by sample population. Thus, this pilot study provides first step towards developing a koala-specific allostatic load index based on multiple stress biomarkers to understand chronic stress in rescued koalas.

**Lay summary:** Stress in koalas can be challenging for their welfare and conservation. In this study, we tested plasma glucocorticoids and their ratios as biomarkers of acute and chronic stress. Our finding showed ratios of DHEAS and DHT to cortisol are comparable across stress parameters and animal demographic characteristics. This study serves as a foundational framework for developing a stress index based on multiple biomarkers that could be useful tool for koala welfare.

## Introduction

Koalas (*Phascolarctos cinereus*) are one of the most iconic marsupial species endemic to Australia, with a specialist folivorous diet in Eucalyptus along the eastern coast of Australia, including Queensland, New South Wales (NSW) and Australian Capital Territory (ACT) (Melzer *et al*., 2000). However, the koala population has experienced a significant decline since the 1990s and has been listed as “threatened” by the Australian Federal Environment and Biodiversity Protection Act 2012 (Gonzalez-Astudillo *et al*., 2017) and most recently its status has been updated to Endangered in South-east Queensland. Koala survivorship is heavily dependent on habitat due to its specialist diet (Smith *et al*., 2013), thus threatened by habitat loss due to rapid land clearing, urbanisation (Gonzalez-Astudillo, *et al.*, 2017, Schipper *et al*., 2008), and increasing frequency and severity of extreme weather events due to climate change (Wallis, 2013). As a result of habitat loss, koalas are often forced to disperse outside of their natural distribution and subsequently experience more anthropogenic threats, including dog attacks and vehicle collisions ([DEWHA]2009, Obendorf, 1983). Another critical threat to the koalas is disease, mainly infection with Chlamydia spp. (*C. pneumoniae* and *C. pecorum*) and koala retrovirus (Gonzalez-Astudillo, *et al.*, 2017). It was found in a 29-year analysis that more than half of the diseased koalas admitted to rescue centres showed signs of Chlamydia (Charalambous *et al*., 2020). Furthermore, low genetic diversity and inbreeding depression in koalas further increase vulnerability to environmental changes and diseases (Houlden *et al*., 1996, Tsangaras *et al*., 2012). Diseases and the other aforementioned threatening factors not only impact koala survivorship directly but also affect their psychological well-being as acute or chronic stressors, especially for rescued koalas with impaired health conditions. Moreover, stress response is very costly in terms of energy (Selye, 1973), exacerbating the welfare of koalas with limited resources of energy due to their low-energy diet (Larsen *et al*., 2014).

Stress in animals is an imbalance in hormones as a result of the animals’ response to cope with stimuli (or stressors) that disrupt their homeostasis (Sheriff *et al*., 2011). The process of animals coping with stress to restore homeostasis is called allostasis (Dickens *et al*., 2010, Romero *et al*., 2019), and allostatic load is the accumulation of stress when a stressor persists (Edes *et al*., 2018). The hypothalamic–pituitary–adrenal (HPA) axis will be activated to release glucocorticoids (GCs) when animals are exposed to stressors (Ralph *et al*., 2016, Wasser *et al*., 2000), for example, the acute and chronic stressors mentioned above for koalas. The HPA axis consists of the hypothalamic paraventricular nucleus (PVN), the anterior pituitary gland, and the adrenal cortex (Sheriff, *et al.*, 2011). Under a stress response, the activated PVN will release neurons to regulate the secretion of corticotrophin-releasing hormone (CRH) (Sheng *et al*., 2021). The CRH then acts on the anterior pituitary to release adrenocorticotropin hormone (ACTH) into the bloodstream (Sheriff, *et al.*, 2011), which then stimulates the adrenal cortex to release GCs (cortisol or corticosterone (CORT)) (Sheng, *et al.*, 2021, Sheriff, *et al.*, 2011). A rise in the GC level allows the animals to mobilise energy by suppressing and changing their essential life-history functions to cope with changes (Bayazit, 2009, Rich *et al*., 2005, Sheriff, *et al.*, 2011, Wingfield *et al*., 2015). This rise in the GC level in response to acute stressors can last from minutes to hours until regulated by a negative feedback mechanism to return to the basal level (Sheriff, *et al.*, 2011, Wingfield *et al*., 2011). However, the HPA axis and the stress response remains activated under chronic stress conditions (Sheriff, *et al.*, 2011). Prolonged activation of the HPA axis leads to allostatic load and has detrimental effects on the animals, including reducing reproductive performances (Whirledge *et al*., 2010), suppressing growth and development (Narayan *et al*., 2016, Whirledge, *et al.*, 2010, Young *et al*., 2004), suppressing the immune system (Acevedo-Whitehouse *et al*., 2009, O’connor *et al*., 2000) which further contribute to the increased vulnerability to disease and infections (Narayan & Williams, 2016; Quan et al., 2001), and affecting the animals’ survivorship and wellbeing in general (McEwen *et al*., 2003). Cortisol is the primary GC found in marsupials (Sheriff, *et al.*, 2011) and can be measured in biological samples such as plasma (Bayazit, 2009, Sheriff, *et al.*, 2011). Measuring the level of cortisol is adopted by most studies on animal stress and endocrinology. Plasma cortisol level is better at detecting acute stress by providing a point-in-time snapshot of the animals’ endocrine activity (Whitham *et al*., 2020). However, cortisol will gradually decline to basal level even when the stressor persists and fails to detect chronic stress in animals. For example, Gundlach *et al*. (2018) failed to detect differences between the cortisol levels of healthy and diseased seals. As a result, using cortisol could solely provide information on acute stress but failed to detect chronic stress and understand the homeostasis profile of the animals.

Nevertheless, the ratio between cortisol and dehydroepiandrosterone (DHEA) was found to be a useful biomarker for assessing chronic stress (Gundlach, *et al.*, 2018, Kamin *et al*., 2017, Longcope, 1996). DHEA and its more stable sulphate ester DHEAS (collectively referred to as DHEA[S]) are adrenal androgen secreted under stress in response to ACTH (Dutheil *et al*., 2021, Nguyen *et al*., 2008). DHEAS shows no strong diurnal rhythm and day-to-day variation due to a longer half-life and a lower rate of metabolic clearance (Kamin, *et al.*, 2017, Longcope, 1996), making it a better biomarker to detect chronic stress (Longcope, 1996). Moreover, DHEA[S] have an antagonising effect on cortisol, i.e., as cortisol level increases, DHEA[S] level decreases, making the ratio between them a useful tool to reflect the endocrine activity (Gundlach, *et al.*, 2018, Kamin, *et al.*, 2017). Both hormones increase at the beginning of the stress response (Dutheil, *et al.*, 2021), however, cortisol increases or remains unchanged and DHEAS declines when the stressor becomes chronic (Whitham, *et al.*, 2020). This difference enables the ratio between them to detect chronic stress. Furthermore, another androgen dihydrotestosterone (DHT), a 5α-reduction androgen of testosterone (T), also has an important role in regulating stress reactivity. During a stress response, the DHT level will be reduced to increase cortisol and ACTH levels (Handa *et al*., 2009, Wingfield *et al*., 1982). DHT’s inhibitory effect on cortisol makes the ratio between them a potential biomarker to measure stress in animals.

Furthermore, cortisol, DHEAS, and DHT could all be measured in blood using EIA (Dutheil, *et al.*, 2021, Monti *et al*., 2012, Ootake *et al*., 2021, Pieper *et al*., 2000, Rosado *et al*., 2010), making it convenient for assessing rescued koalas that must take blood tests at admissions. To our knowledge, there is no study on using DHEAS, DHT, and the ratio between cortisol and them to detect stress in koalas. Therefore, this study aims to determine if the ratio of cortisol to DHEAS and DHT are better tools for assessing chronic stress in rescued koalas than plasma cortisol alone.

## Methods

### Ethics approvals

All animal handling procedures were approved by the Animal Ethics Committee (Native and Exotic Wildlife and Marine Animal Group) at the University of Queensland and The Royal Society for the Prevention of Cruelty to Animals (RSPCA Queensland) (hereafter referred to as RSPCA) (project number: 2023/AE000750).

### Study animals and sampling design

This study included 10 rescued koalas (F = 3, M = 7) admitted to RSPCA. A 1.5 mL of whole blood sample was collected from each koala at the initial health inspection shortly after arrival by the RSPCA veterinarians. A 2cm x 2 cm section of fur covering the cephalic vein was shaved, and the blood was collected under anesthesia with a 3 ml syringe and 23-g, 5/8-inch needle. The blood was stored in EDTA tubes, labelled with individual koala ID, and froze immediately in the freezer until collected. The rescued koalas remained at RSPCA for diagnosis and treatment until they were released, euthanised, or transferred to other facilities. Samples were transported to the laboratory in a chiller bag with at least one ice bag and kept in the freezer for less than a month until assays.

Furthermore, the following information was also recorded during the initial health inspection: koala ID, name, gender, age, weight, body conditioning score (BCS), and reason for admission. The age of the koalas was determined based on the dentition, primarily the progressive stage of wearing of the upper premolar and first molar teeth (Blanshard *et al*., 2008), during anesthesia in the initial health inspection. BCS was also measured at the initial health inspection based on Blanshard, *et al.* (2008). The assessment was done *via* palpation of muscle mass over the scapulae by placing hands across the koalas’ shoulders and rubbing over the muscles (Blanshard, *et al.*, 2008). The BCS was recorded based on the numerical body condition scoring system between 1 and 5, where 5 is excellent, 4 is good, 3 is fair, 2 is poor, and 1 is emaciated., and rescaled to a scale of 10.

### Laboratory validation

The laboratory validation of all assays was based on Narayan *et al*. (2013). A parallelism between serial dilutions of pooled hormones and standard curves was done to check the parallel displacement of the hormone standard against the pooled extracts (Figure 2-4). The hormone concentration data was log-transformed for all tests. The standard curves were displayed as linear regression equations: y = mx + b. In this equation, y is the relative optical density (OD), x is the hormone concentration, b is the y-intercept, and m is the slope of the linear regression line.

**Figure 1.**
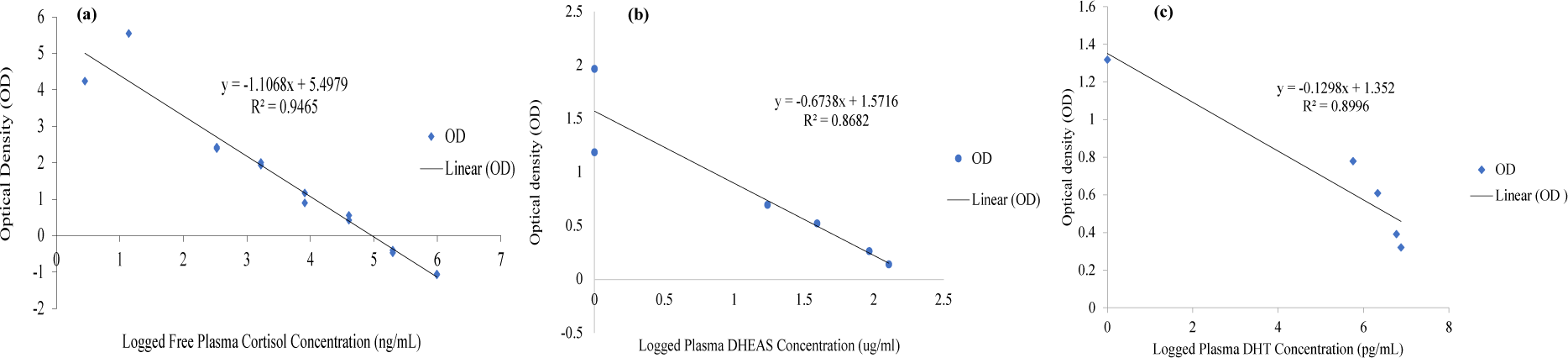
The parallelisms between serial dilutions of pooled cortisol (a), dehydroepiandrosterone sulphate (DHEAS) (b), and dihydrotestosterone (DHT) (c). The standard curve is displayed as a linear regression equation: y = mx + b. Y is the relative optical density (OD), x is the hormone concentration, b is the y-intercept, and m is the slope of the linear regression line. The R² calculates the recovery of hormones as per the regression equation by multiplying R² by 100. The assay sensitivity for cortisol EIA was 9.14 pg/well, and the coefficient of variation was 5.5%. For DHEAS, the assay sensitivity was 3.45 ug/mL and the coefficient of variation was 3.97%. For DHT, the assay sensitivity was 0.32 ng/mL and the coefficient of variation was 12.04%.

**Figure 2.**
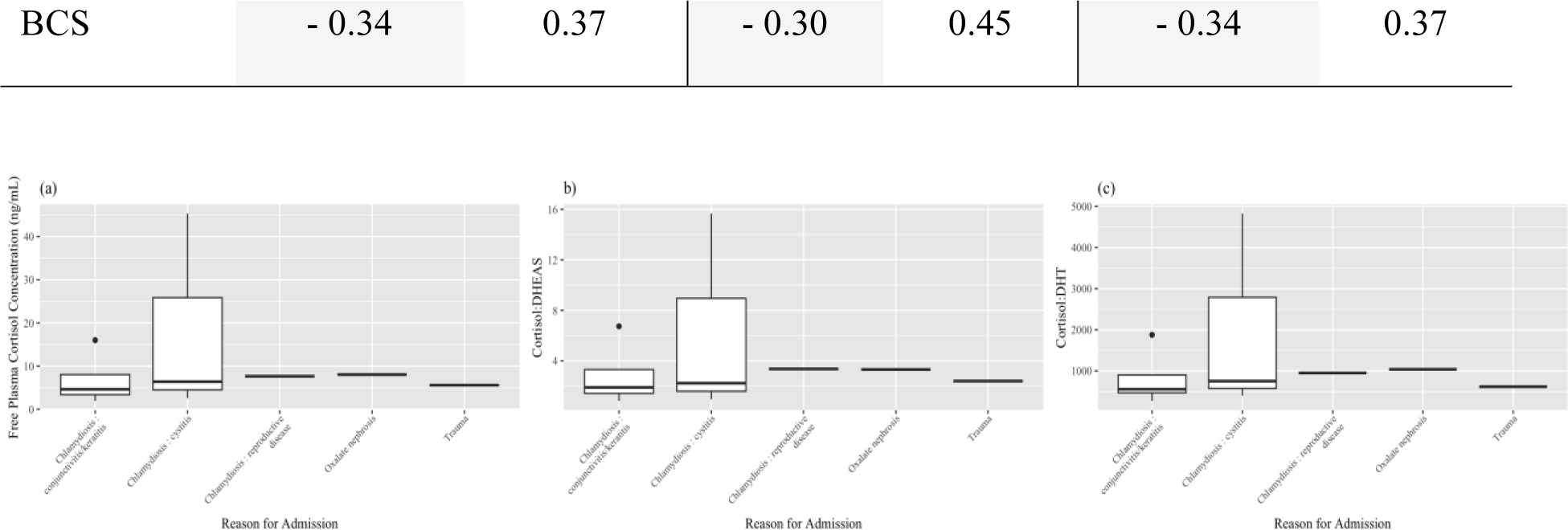
The distribution of three plasma biomarkers: (a) free plasma cortisol concentration, (b) cortisol:dehydroepiandrosterone sulphate, and (c) cortisol:dihydrotestosterone (DHT) across different reasons for admission of rescued koalas at RSPCA Queensland (n = 10). The median is indicated by the bold line.

### Free plasma cortisol EIA

The free plasma cortisol EIA was done following guidelines provided by Brown *et al*. (2004) and Narayan, *et al.* (2013). In this EIA, a polyclonal anticortisol antiserum (R4866) diluted 1:15,000 with a 100% cross-reactivity with cortisol (Munro and Stabenfeldt, 1985, Narayan et al., 2010ined), horseradish peroxidase conjugated cortisol label diluted 1:80,000, and standards were used to determine the concentration. A Nunc MaxiSorp 96-well plate was used in this EIA. The plate was first coated with 50 μL of cortisol antibody diluted to the appropriate concentration using coating buffer and then incubated for at least twelve hours at 4 °C. After incubation, the plate was washed with an automated plate washer with phosphate-buffered saline containing 0.5 ml L^−1^ Tween 20 to remove unbound material. Ten stocks of standards, controls, and samples were diluted in an assay buffer, and 50 μL of each was added to respective wells. Each sample was replicated twice, and each standard was replicated three times. For each well, 50 μL of the corresponding horseradish peroxidase label was added and incubated at room temperature for two hours. After incubation, the plate was washed with the same wash solution, and 50 μL of a substrate buffer (0.01% tetramethylbenzidine and 0.004% H_2_O_2_ in 0.1 M acetate citrate acid buffer, pH 6.0) was added to all wells. The plate was then incubated at room temperature. After 7 to 10 minutes, the zero well reached an optical density (OD) of 0.7-1.0 based on visual inspection. The reaction was stopped by adding 50 μL of stop solution to each well. The plate was then read at 450 nm (correction 630nm) on an EL800 (Bio Tek) microplate reader.

### Plasma DHEAS enzyme-linked immunosorbent assay (ELISA)

An ELISA Kit (product code ab108669; from Abcam, Australia; additional information available at https://www.abcam.com/en-au/products/elisa-kits/dhea-sulfate-dhea-s-elisa-kit-ab108669<u>#</u>) was used for processing and analysing the concentration of plasma DHEAS. For this ELISA, a 96-well plate precoated with anti-DHEA sulphate antibodies is used. The samples were diluted to the appropriate concentration in serum diluent. 30 μL of five stocks of standards, controls, and diluted samples were added to the respective wells, and each was replicated twice. For each well, 100 μL of DHEA sulfate-HRP conjugate was then added with an empty well left as the substrate blank without the conjugate. The plate was then covered with foil and incubated for one hour at 37°C. After incubation, the plate was washed three times using an automated plate washer with 300 μL diluted wash solution to remove unbound material. For all wells, 100 μL of TMB substrate solution was added and incubated for exactly fifteen minutes at room temperature in the dark. The reaction was then stopped by adding 100 μL stop solution into all wells, and the plate was gently agitated using a rotating mixer. The plate was then read at 450 nm on an EL800 (Bio Tek) microplate reader.

### Plasma DHT ELISA

An ELISA Kit (product code ab287824; from Abcam, Australia; additional information available at (https://www.abcam.com/ab287824) was used for processing and analysing the concentration of plasma DHT. The lyophilised DHT standard was reconstituted to prepare the standard by adding 1 ml of standard dilution Buffer to make a stock solution. A 0.6 ml of 1250 pg/ml top standard was then made by adding 0.3 ml of the stock solution in 0.3 ml of the standard/sample dilution buffer. A 2-fold serial dilution was then performed to make the standard curve within the range of this assay. The plate was washed twice using an automated plate washer with 1X wash solution first, and 50 μl of prepared standards and samples were then added into respective wells with one replicate. 50 μl of biotin-labelled antibody working solution diluted with antibody dilution buffer was then added immediately into each well. The plate was then covered with a plate sealer and incubated for 45 minutes at 37°C. After incubation, the solution was discarded and the plate was washed three times with 350 μl of wash solution. For each well, 0.1 ml of HRP-Streptavidin Conjugate (SABC) working solution diluted with SABC dilution buffer was added. The plate was then covered and incubated for 30 minutes at 37°C. After incubation, the solution was discarded and washed five times with 350 μl of wash solution. 90 μl of TMB substrate was then added into each well, and the plate was covered and incubated for 15 minutes at 37°C in the dark. The reaction was stopped by adding 50 μl of the stop solution. The plate was then read at 450 nm on a EL800 (Bio Tek) microplate reader.

### Statistical analysis

Statistical analyses were performed using R Studio (version 4.3.3; R Core Team 2024) and Microsoft® Excel. Free plasma cortisol, DHEAS, and DHT concentration were calculated based on the standard curve and OD. Three biomarkers, including the free plasma cortisol, cortisol:DHEAS ratio, and cortisol:DHT ratio, were assessed against five factors, including the reason for admission, gender, age, weight, and BCS. The alpha level for detecting the significant differences was 0.05. Data are presented as median ± standard error (SE). The reasons for admission factor have five levels, including Chlamydiosis: cystitis, Chlamydiosis: conjunctivitis/keratitis, Chlamydiosis: reproductive disease, oxalate nephrosis, and trauma. Kruskal-Wallis rank sum tests were conducted to determine the statistical difference between different reasons for admission for all biomarkers because they did not follow normal distributions. The missing values in the data were listed as non-applicable (NA) and dealt with by omitting rows with NA values but including NA in the same position as the omitted observations in the input data to maintain the alignment. The biomarkers were plotted against reasons for admissions and displayed as boxplots to show distributions. The Wilcoxon rank-sum tests were conducted to detect the significant differences in the median of three biomarkers between different genders and age groups because they failed to follow a normal distribution. The biomarkers were plotted against gender groups (F/M) and age groups and displayed as bar plots. For this test, koalas were assigned to the following age groups based on their life history (Charalambous, *et al.*, 2020).

1. Joey: koala between the ages of birth to 6 months
2. Juvenile: koala aged between 6 months and 1 year
3. Adult: koala aged between 1 year and 7 years
4. Old: koala aged over 7 years

Spearman’s rank correlation coefficients (r) were calculated for age (numerical without grouping), weight, and BCS to determine the relationship between the factors and the biomarkers. The strength of the relationships was determined based on the absolute value of r and classified into 5 strengths, including a negligible correlation (0 < |r| ≤ 0.10), a weak correlation (0.10 < |r| < 0.39), a moderate correlation (0.40 < |r| < 0.69), a strong correlation (0.70 < |r| < 0.89), and a very strong correlation (0.9 < |r| < 1.0) (Schober *et al*., 2018). The biomarkers were plotted against age, weight, and BCS and displayed as scatterplots.

## Results

### Study animals and sampling design

The general data of investigated rescued koalas were collected and collated in Table 1.

**Table 1.**
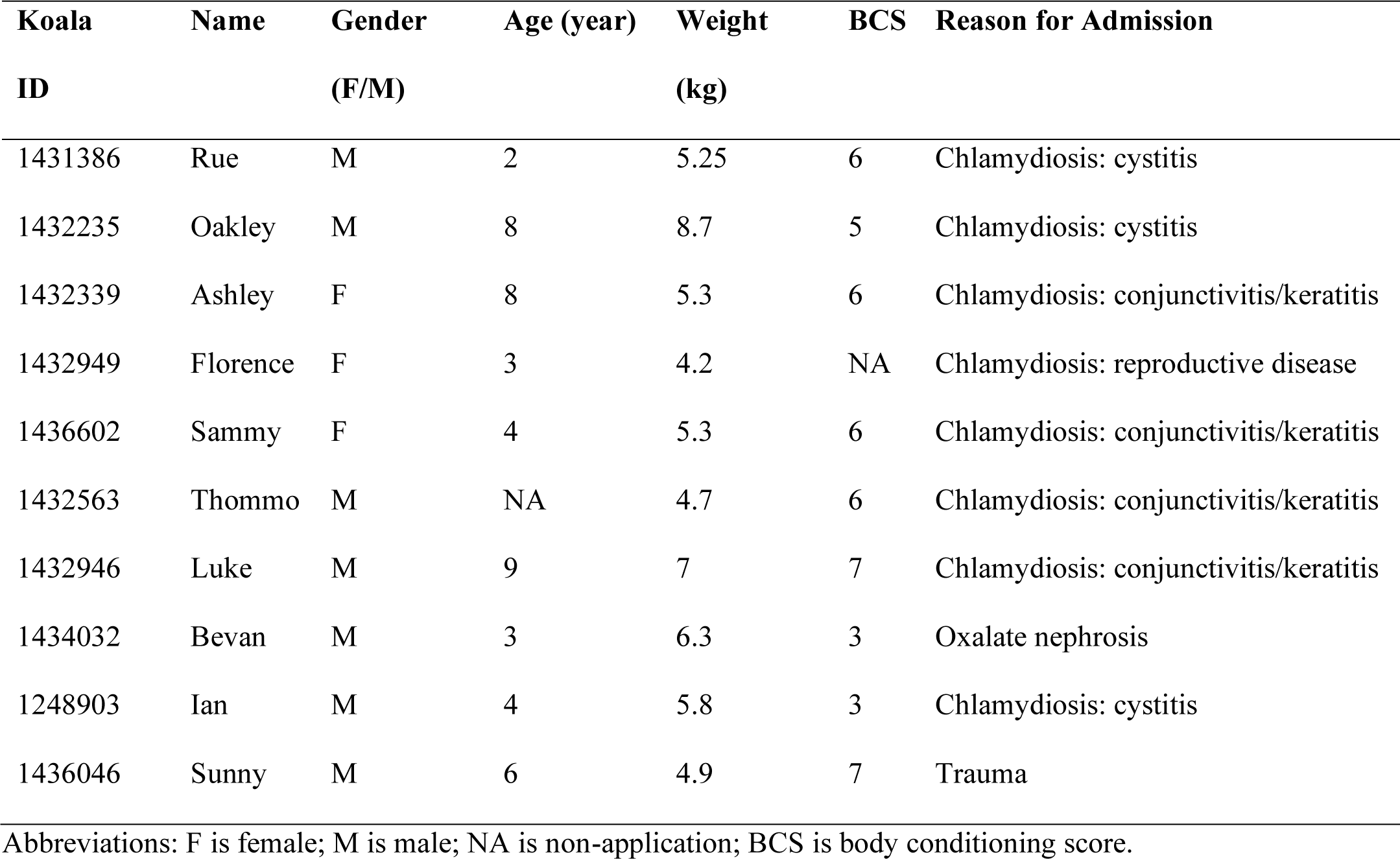
Summary of the general data investigated in rescued koalas (n = 10) at RSPCA Queensland.

### Laboratory Validation

The regression equation of each hormone based on parallelism results was as follows (Figure 1a-c):

1. Cortisol: y = -1.1068x + 5.4979, R² = 0.9465
2. DHEAS: y = -0.6738x + 1.5716, R² = 0.8682
3. DHT: y = -0.1298x + 1.352, R² = 0.8996

Therefore, the accuracy of displacement of hormones as per the regression equation were 94.7%, 86.8%, and 90.0%, respectively.

### Comparison between biomarkers across factors

#### Reason for admissions

There was no significant difference in the median of free plasma cortisol concentration, cortisol:DHEAS ratio, and cortisol:DHT ratio between different reasons for admission (p = 0.79, 0.80, 0.79, respectively). Among all reasons for admissions, Chlamydiosis: reproductive disease and oxalate nephrosis tend to have the highest medians (Table 2). Oxalate nephrosis had the highest median in free plasma cortisol concentration and cortisol:DHT, while Chlamydiosis: reproductive disease had the highest median in cortisol:DHEAS (Table 2). In addition, Chlamydiosis: conjunctivitis/keratitis had the lowest medians across all biomarkers (Table 2). The distribution is similar based on visual inspections across biomarkers with outliers affecting the Chlamydiosis: conjunctivitis/keratitis and the Chlamydiosis: cystitis (Figure 2).

**Table 2.**
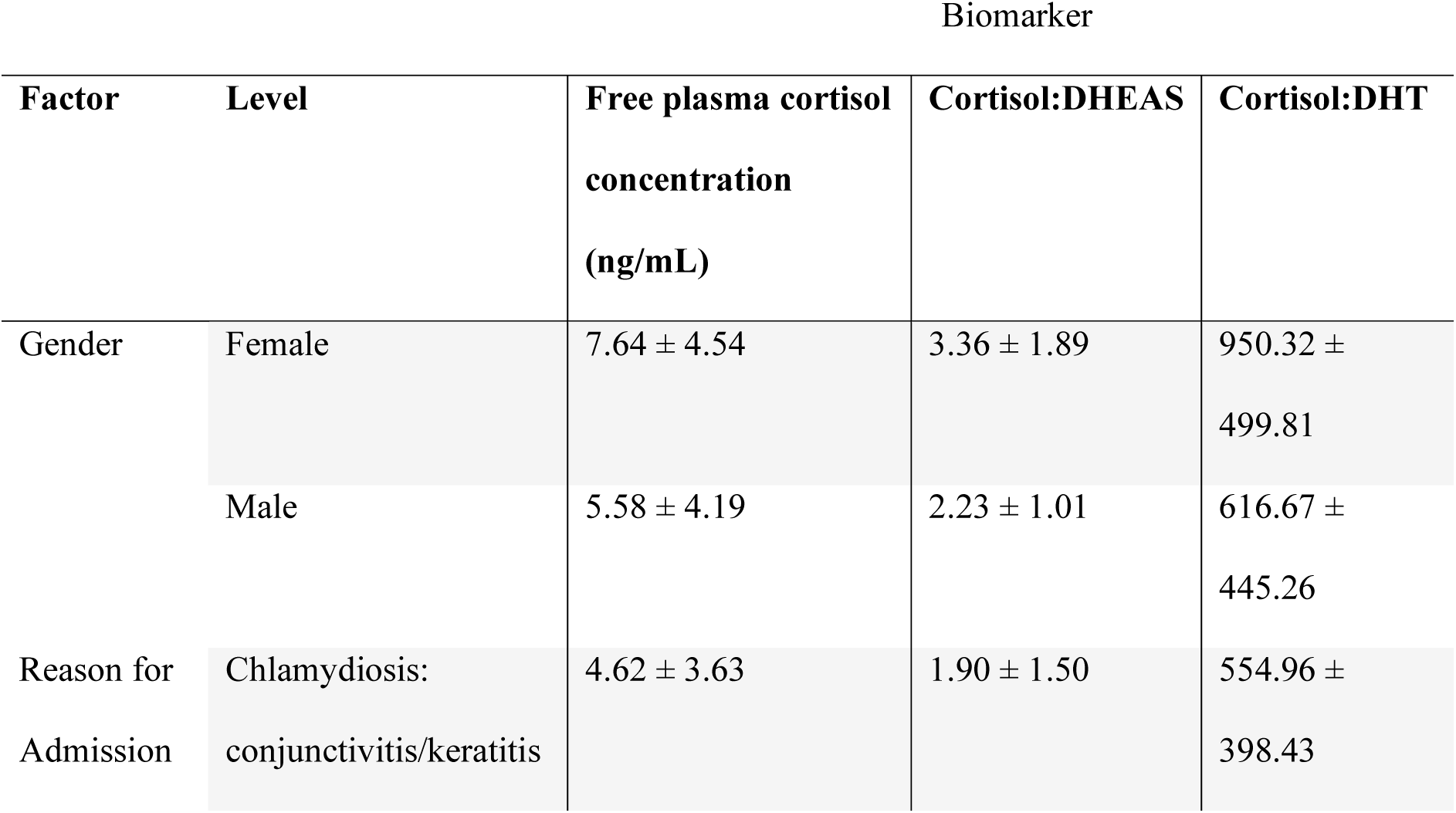
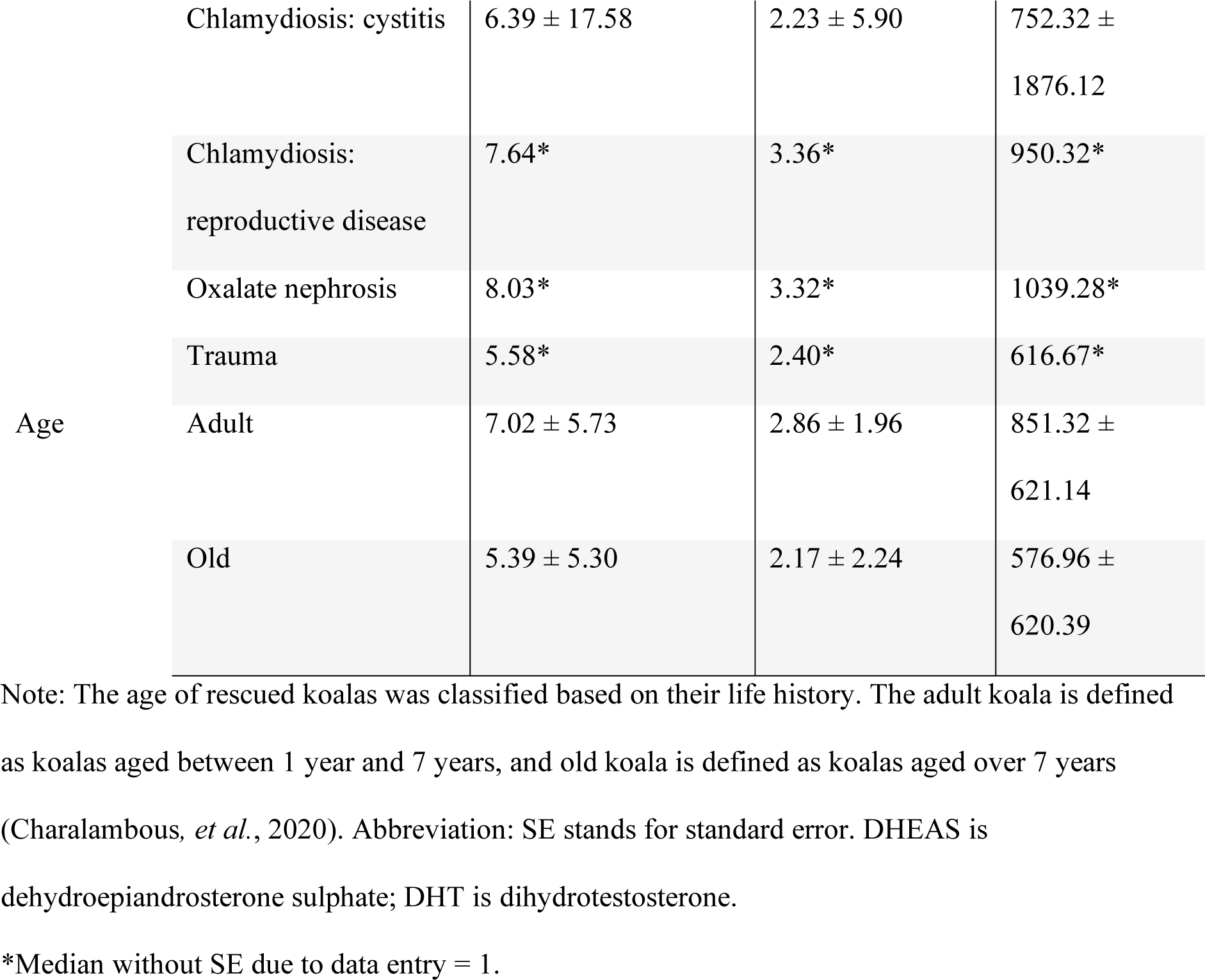
The endocrinological results of three biomarkers: free plasma cortisol concentration, cortisol:DHEAS, and cortisol:DHT of rescued koalas (n = 10) at RSPCA Queensland. Data is presented as median ± SE.

#### Gender

There was no significant difference in the median of free plasma cortisol concentration, cortisol:DHEAS ratio, and cortisol:DHT ratio between female and male recused koalas (p = 0.67, 0.52, 0.67, respectively). However, females have a higher median than males in general (Table 2; Figure 3).

**Figure 3.**
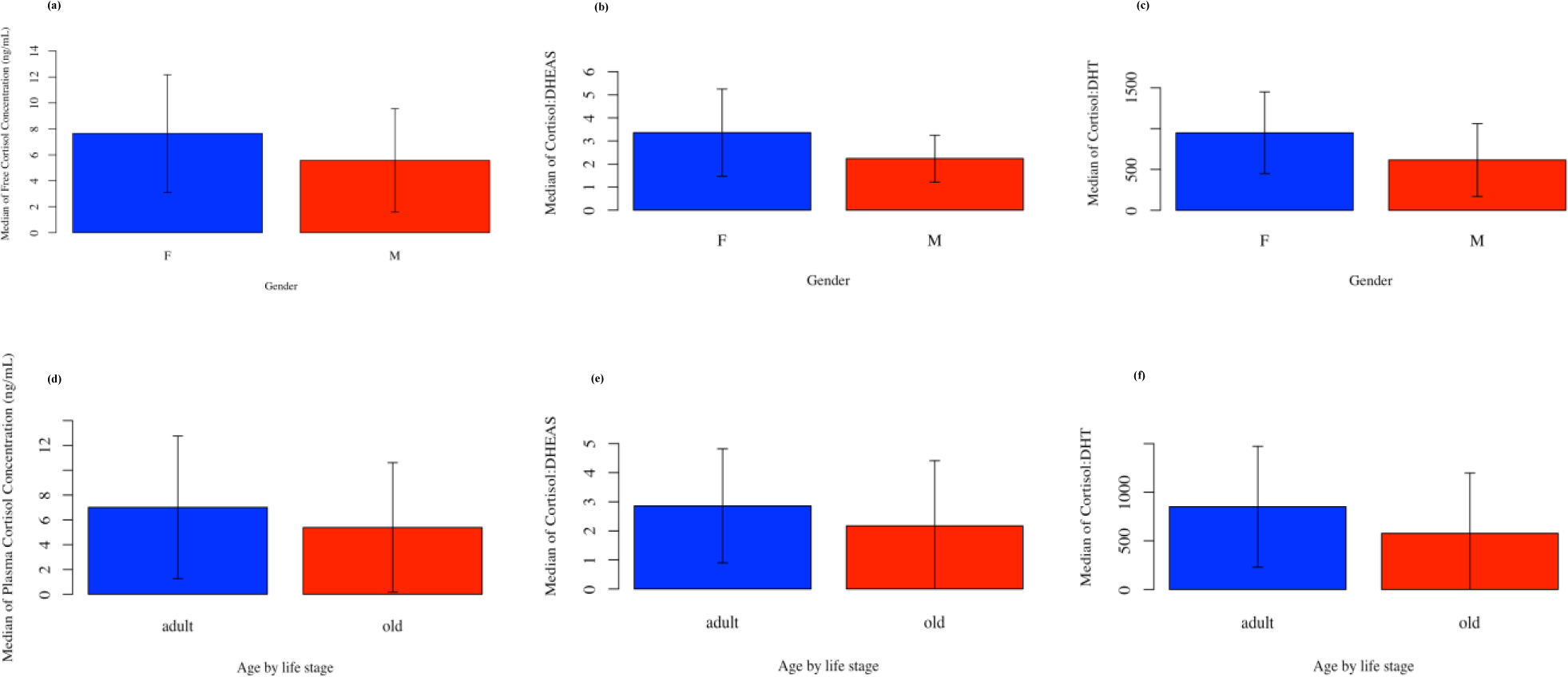
The comparison of the median of three biomarkers: (a) free plasma cortisol concentration, (b) cortisol:dehydroepiandrosterone sulphate, and (c) cortisol:dihydrotestosterone (DHT) showing that female (F) always has a higher median than male (M); and the comparison of the median of three biomarkers (c) free plasma cortisol concentration, (d) cortisol:dehydroepiandrosterone sulphate, and (e) cortisol:dihydrotestosterone (DHT) showing that the adult koalas always has a higher median than the old koalas at RSPCA Queensland (n = 10). The standard errors are presented as the error bars. The F stands for the female and M stands. The adult koala is defined as koalas aged between 1 year and 7 years, and old koala is defined as koalas aged over 7 years (Charalambous, *et al.*, 2020). The standard errors are presented as the error bars.

#### Age by life stage

The adult and the old rescued koalas displayed no significant difference across the three biomarkers (p = 0.55, 0.55, 0. 55, respectively). Although insignificant, adult koalas have a higher median than the old koalas in general (Table 2; Figure 3).

#### Age, weight, and BCS

All the correlations between factors and biomarkers were insignificant (Table 3). All the correlations between age and BCS and biomarkers were negative (Table 3). On the contrary, all correlations between weight and biomarkers were positive except for cortisol:DHEAS (r = - 0.08) (Table 3). The strength of relationships between free plasma cortisol concentration and age, weight, and BCS is weak, negligible, and weak, respectively (Table 3). For cortisol:DHEAS, their strength of relationships with age, weight, and BCS is weak, negligible, and weak, respectively (Table 3). For cortisol:DHT, their strength of relationships with age, weight, and BCS is weak, negligible, and weak, respectively (Table 3).

**Table 3.**
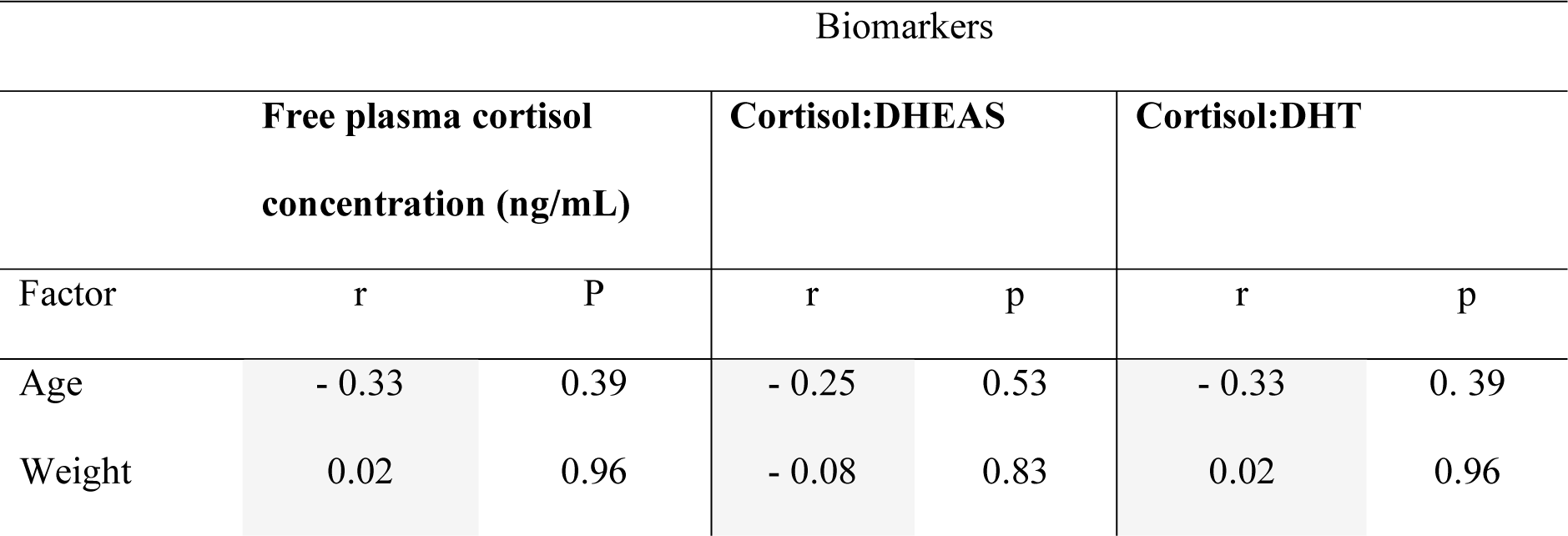
The Spearman’s rank correlation between the age, weight, body conditioning score (BCS) and three biomarkers including free plasma cortisol concentration, cortisol: dehydroepiandrosterone sulphate, and cortisol:dihydrotestosterone (DHT) of the rescued koalas at RSPCA Queensland (n = 10). The correlation coefficient is presented as r, and p-value is presented as p.

## Discussion

This study investigated the use of three biomarkers, free cortisol concentration, cortisol:DHEAS, and cortisol:DHT, to detect chronic stress in rescued koalas by comparing the levels of each biomarker in different groups of koalas. In summary, no significant differences were found between all biomarkers across different groups of koalas. Nevertheless, we identified consistent trends across biomarkers to assess chronic stressors, including Chlamydiosis: reproductive disease, oxalate nephrosis, and conjunctivitis/keratitis. Consistency across biomarkers was also found between gender, age, weight, and BCS, implying the potential use of these biomarkers to assess stress together.

### Reason for admission

#### Chlamydiosis: reproductive disease and Oxalate nephrosis

Disease caused by *C. pecorum* is one of the reasons for admission included in this study. *C. pecorum* is the most pathogenic Chlamydia species, leading to ocular and urogenital diseases (Fabijan *et al*., 2019). It is the only species included in this study, probably due to a higher prevalence in wild koalas than *C. pneumoniae*, which is more common in captive populations (Blanshard, *et al.*, 2008). Out of all Chlamydia diseases, reproductive diseases have the highest median in cortisol:DHEAS and the second highest median in the other two biomarkers in our study. Chlamydia infection could establish in the female and male reproductive tract, leading to inflammation, fibrosis, scarring and infertility (Blanshard, *et al.*, 2008, Quigley *et al*., 2020). It is more difficult to detect acute reproductive diseases due to the lack of evident external clinical signs except for possible mucopurulent exudate from the opening (Polkinghorne *et al*., 2013a). Chronic conditions are more evident in females with cystic changes (Polkinghorne, *et al.*, 2013a) but less evident in males due to uncommon sonographic changes (Loader, 2010). It is difficult to detect moderate to severe inflammation of the reproductive tract without the presence of cystitis, but the animals could be under severe underlying pain and disease (Hemsley *et al*., 1997). The koalas included in this study were all admitted to RSPCA due to the presence of obvious external clinical signs. Therefore, the koalas with reproductive diseases in our study probably suffer from chronic Chlamydiosis and experience chronic pain and stress. Under chronic stress, a higher level of plasma cortisol is expected due to dysregulation of the HPA axis and prolonged elevation of cortisol (Sapolsky, 1990). This is also supported by Narayan *et al*. (2012), who found a higher cortisol concentration in koalas suffering from chronic stress and ongoing health issues. Therefore, although insignificant, our result showed that all three biomarkers have the potential to detect chronic stressors.

Another reason for admission with high medians across three biomarkers is oxalate nephrosis. Oxalate nephrosis is one of the most prevalent diseases in koalas from South Australia (Speight *et al*., 2012). Oxalate nephrosis is characterised by the presence of calcium oxalate deposits in the tubules of the kidneys (Speight *et al*., 2019). This disease is progressive (Speight *et al*., 2013) and is considered relatively chronic because it is associated with renal dysfunction caused by inflammation, tubule obstruction, and fibrosis, eventually leading to renal failure (Speight, *et al.*, 2019). Furthermore, koalas with oxalate nephrosis showed clinical signs of renal dysfunction, including weight loss, polydipsia, polyuria (Haynes *et al*., 2004), loss in body condition (Speight, *et al.*, 2013), and azotaemia (Speight, *et al.*, 2013). Affected koalas also experience dehydration and increased thirst due to kidney dysfunction and inability to conserve water (Haynes, *et al.*, 2004). These could all contribute to chronic pain and illness in koalas and, as a result, koalas with oxalate nephrosis often require euthanasia due to welfare concerns (Speight *et al*., 2020). Furthermore, the cause of oxalate nephrosis is unclear and suggested to be very complex and multifactorial (Narayan, *et al.*, 2016).

Except for dietary intake of oxalate (Narayan, *et al.*, 2016), possible stressors that could lead to oxalate nephrosis are chronic and environmental, including heat stress, car and dog impacts, maternal stress, nutritional deprivation, and dehydration (Narayan, *et al.*, 2016). Therefore, koala with oxalate nephrosis was likely to be under chronic stress, and the disease further leads to chronic discomfort.

The high median plasma cortisol concentration found in our study is supported by the chronic illness associated with oxalate nephrosis and the expected higher cortisol concentration under chronic conditions (Narayan, *et al.*, 2012, Sapolsky, 1990). The use of cortisol:DHEAS and cortisol:DHT to detect chronic stress is not validated in this study due to the insignificant differences. However, the consistent trend found in reproductive disease and oxalate nephrosis supported their potential usefulness in assessing chronic stressors. One possible limitation contributing to the insignificant result is the small sample size (n = 1) of both diseases included in our study. Another limitation is the stress induced by capture. The median plasma cortisol concentration for reproductive disease and oxalate nephrosis in our study is 7.64 ng/mL and 8.03 ng/mL (Table 3), which is similar to the mean plasma cortisol concentration right after capture compared to 6 hours and 24 hours after capture (7.92 ± 8.36 ng/mL, 3.16 ± 2.6 ng/mL, and 4.52 ± 4.96 ng/mL, respectively) (Speight *et al*., 2016). This suggests that our result may be affected by the capturing process. Such human intervention could influence stress hormones in captivity and increase cortisol concentration (Narayan *et al*., 2011, Ram *et al*., 2005), especially when our sample population is wild without habituation to humans.

#### Conjunctivitis/keratitis

Across all reasons of admission, Chlamydiosis: conjunctivitis/keratitis had the lowest medians across all biomarkers. Conjunctivitis/keratitis due to Chlamydiosis happens when the conjunctiva of the eye is infected (unilateral or bilateral) (Quigley, *et al.*, 2020), leading to inflammation, discharge, conjunctival hyperplasia and fibrosis in acute conditions. As the infection develops to be chronic, blindness could happen due to conjunctival hyperplasia and pathological changes affecting the cornea (Polkinghorne, *et al.*, 2013a). Our median (4.62 ± 3.43 ng/mL) (Table 3) is lower than the baseline plasma cortisol concentration of wild koalas (7.0 ng/ml) (Hajduk *et al*., 1992), suggesting it is abnormal due to either acute or chronic stress conditions. Although conjunctivitis/keratitis is evident and easy to identify in both acute and chronic conditions (Polkinghorne *et al*., 2013b), the low cortisol concentration found in our study is highly possibly the result of a chronic stressor. Nyari *et al*. (2017) proved that infection from mother to joey is a common exposure route of ocular infection related to Chlamydia, resulting in higher chances of chronic infection developed since young. Another study by Cockram *et al*. (1981) also suggested that koalas infected at a young age usually carry and develop the infection and grow to maturity due to the typically long duration of the disease. Different to the elevated stress level observed in previous findings, this lower stress level could be due to habituation.

A prolonged chronic condition could reduce cortisol concentration due to habituation or exhaustion when the animal gets accustomed to the stress (Konarska *et al*., 1989) as the final stage of a stress response (Selye, 1973). For example, Malayan pangolin (*Manis javanica*) rescued from the wildlife trade was found to have a physiological exhaustion due to chronic stress, leading to a lower level of cortisol than pangolins born and reared in captivity (Yan *et al*., 2021). Except for habituation and exhaustion, a low cortisol level could also be the result of a regulated change in the HPA axis.

European starlings (*Sturnus vulgaris*) had a lower CORT level under chronic stress when the hypothalamus regulates the release of arginine vasotocin instead of ACTH, leading to a reduction in ACTH and consequently a lower level of CORT (Rich, *et al.*, 2005). One limitation of our study is the lack of healthy koalas because the sample population is rescued koalas. It is difficult to understand the cortisol response to different stressors without baseline data to compare with (Narayan, *et al.*, 2013). However, the consistent trends observed across biomarkers allowed possible comparisons and demonstrated the potential of all three biomarkers to detect chronic stress. Nevertheless, baseline data is required to further differentiate scenarios with a higher cortisol concentration for reproductive disease and oxalate nephrosis and a lower cortisol concentration for conjunctivitis/keratitis.

#### Gender

Females had a higher median across all biomarkers than males in our study (P > 0.05). Narayan (2019) also found a higher faecal cortisol metabolite level in females in a population of healthy koalas, suggesting that females had a higher level regardless of chronic stress due to a higher metabolism. On the contrary, Webster *et al*. (2017) found a higher faecal cortisol metabolite (FCM) in males exposed to intensive visitor interaction and reported a delayed response in females to intensive visitor interaction. However, Webster, *et al.* (2017) was conducted when koalas were housed with different sexes, leading to a potential impact of social dominance relationships and reproductive status on their result. Their previous study also found that females did not elevate FCM levels unless lactating (Narayan, *et al.*, 2013), suggesting that lactation could impact the cortisol excretion of female koalas. Sex is one of the factors that impact the variation of GC levels the most (Narayan, *et al.*, 2013). Many factors could contribute to the differences between sex-related GC. For example, Goymann (2012) suggested that steroid metabolism and pituitary responsiveness are underlying factors leading to the difference between faecal cortisol metabolites between sexes. Reproductive activity is another factor that could influence sex-related cortisol concentration. Koalas are seasonal breeders, and the breeding season for females is generally between August and April in Queensland (O’Callaghan *et al*., 1991). On the other hand, males could mate at any time but were more active during the breeding season (Blanshard, *et al.*, 2008). Our study was conducted between January and February during the breeding season. All koalas in our study reached sexual maturity (Table 1), which is generally around 15 months for females and 24 months for males (Blanshard, *et al.*, 2008), and one of the females had a pouch young. Therefore, the koalas included in this study are reproductively active, and the differences in cortisol concentration could be sex-related. For example, the cyclic fluctuations of oestrogen and progesterone concentrations in females during breeding season are one of the factors that impact cortisol concentration (Palme *et al*., 2005). The increased metabolic demands associated with reproduction could also influence cortisol levels (Touma *et al*., 2005).

Furthermore, the higher cortisol level found in females could be due to their need to increase watchfulness to protect young from predator (Gray, 1987) and aggressive dominant males (Vandenheede *et al*., 1993). Maternal effort of koalas is another stressor that could impact cortisol concentration except reproduction. Female koalas invest heavily in prolonged maternal care by allocating more energy and resources to ensure the growth and survival of their young (Narayan, 2019, Tobey *et al*., 2006). In addition, the plasma T concentration of male koalas was found to have a marked increase from late July to August, and the average concentration lasted until January (Handasyde *et al*., 1991). As a result, our study conducted between January and February should have a lower DHT concentration, leading to a higher cortisol:DHT. However, the median male cortisol:DHT was lower in our result because our study was compared to females instead of males at different times. Our study expected a higher DHT concentration in males because DHT is the 5α-reduction androgen of T. This should result in a lower cortisol:DHT in males, which was observed. Therefore, our result is generally expected despite being insignificant, and one of the limitations is the unbalanced sample size between sexes (F = 3, M = 7).

#### Age

The stress levels of adult and old koalas were statistically similar (p > 0.05). However, adult groups tend to have a higher level compared to older groups across all biomarkers, and this trend was further supported by the negative relationships found between age and stress level. This was expected because age has a significant impact on the koala immune system by influencing gut microbiota and host immune markers, increasing their susceptibility to *C. pecorum* (Chen *et al*., 2023). There is also a higher chance of disease as the koalas get older, but the increases cease as the koalas become old, i.e., there is a higher prevalence of *C. pecorum* infection in adult koalas than in older koalas (Nyari, *et al.*, 2017). The higher susceptibility and higher disease prevalence could be the underlying reasons for the higher stress level observed in adult koalas. Furthermore, evidence shows that most koalas with *C. pecorum* were infected at an early age (Jackson *et al*., 1999), resulting in a longer duration of stress in older koalas. However, koalas in our study all had clinical signs, and differences in disease prevalence could potentially had less impact. Habituation to chronic stressors is one of the key theoretical explanations proposed by Romero (2004) for individual differences in stress hormone concentration, which is another possible explanation for our result due to the longer duration of exposure to chronic stress in older koalas. Dampening of stress response to conserve energy and avoid unnecessary overreaction could happen when the stressor has chronic impacts on fitness because stress response is costly in terms of energy (Bonier *et al*., 2009, Rankin *et al*., 2009). This was also observed in alpine chamois (*Rupicapra rupicapra*) that have habituated to chronic environmental stressors, and further exposure to stressors would not result in further stress (Anderwald *et al*., 2021). Another study of spotted salamanders (*Ambystoma maculatum*) also found a lower GC concentration in the population exposed to chronic environmental stressors than in an undisturbed population (Homan *et al*., 2003). Habituation to chronic stressors was also observed in heavily disturbed birds with a lower CORT level (Arlettaz et al. 2007in51;51). It is also possible that older koalas were in poorer body condition to initiate an efficient stress response due to poor functioning of the HPA axis, leading to a reduced stress hormone concentration (Anderwald, *et al.*, 2021, Romero, 2004, Taillon *et al*., 2008). Furthermore, all koalas were exposed to acute stressors such as capturing and handling in our study, which could lead to an elevated cortisol level. This impact of short-term stressors in older koalas may be overlapped by the prolonged exposure to chronic stressors through habituation (Rich, *et al.*, 2005), leading to a reduced cortisol level in older koalas. In addition, the higher stress level of adult koalas could be due to their higher need to survive and perform life-history functions, including mating (Charalambous, *et al.*, 2020). Furthermore, older animals with lower reproductivity were found to have a reduced stress response compared to mature animals to ensure reproduction still happens (Wilcoxen *et al*., 2011). Therefore, the results of age agreed with each other and were supported by literature. This displayed the possibility of using all biomarkers together to assess stress despite their insignificance.

#### Weight

Weight was the only factor having positive relationships with the biomarkers (p > 0.05). The positive correlations were not expected because we expect a poor health condition and subsequently lower weight as the koala gets more stressed. However, cortisol:DHEAS has a negative correlation with weight and this relationship was supported by literature. This negative relationship could be explained by the fact that koalas with larger bodies need more water (Nagy *et al*., 1985), which may pose chronic stress on koalas to find water resources and result in a lower cortisol level. The disagreement in relationships observed may be due to the difference between chronic and acute stressors and suggests that the cortisol concentration and cortisol:DHT increases as koalas get heavier, which could be due to more acute stressors. Although insignificant, our hypothesis that cortisol:DHEAS is a better biomarker for detecting chronic stress is partially supported. However, the abilities of different biomarkers to detect chronic and acute stressors were invalidated in our study, and the disagreeing relationships observed that are different from age and BCS may be due to limitations. Another explanation for the disagreement is the complex relationship between stress and weight. For example, body size in koalas is sex-related (Tobey, *et al.*, 2006), and the need for water is temperature- and habitat-related (Narayan, *et al.*, 2016). Furthermore, body size is also age-dependent for male koalas due to sexual selection (Briscoe *et al*., 2015, Charlton *et al*., 2012, Tobey, *et al.*, 2006). Blas *et al*. (2006) suggested that age in birds is related to increased weight, which is associated with larger energy demand and results in an elevated stress hormone level to survive (Blas, *et al.*, 2006). Fat reserve is another factor that could potentially affect this relationship due to the stress hormone’s ability to mobilise energy stores (Breuner *et al*., 2008, Long *et al*., 2004, Sapolsky *et al*., 2000). Our result could also be affected by the fact that all koalas in this study suffer from health issues that could impact their weight. Therefore, the relationship between stress and weight is complex and multifactorial, which may explain the inconsistent relationships observed in weight compared to other factors.

#### BCS

We found non-significant negative relationships between BCS and all biomarkers (p > 0.05), suggesting that the stress level increases as the koalas get less healthy. Chronic disease could result in poor body condition not only due to painfulness but also a higher energy demand for recovery (Fuller *et al*., 2011, Tompkins *et al*., 2011). A dampened HPA axis reactivity was also observed as body condition improves (Beale *et al*., 2004, Breuner, *et al.*, 2008). A better body condition allows individuals to not perceive stressors as stressful as unhealthy individuals; for example, a lack of food resources is less stressful for healthier individuals (Le Ninan *et al*., 1988). A negative relationship between stress and BCS was also observed in other species using different biomarkers, including birds (Breuner *et al*., 2003, Cherel *et al*., 1988, Gray *et al*., 1990, Hood *et al*., 1998, Long, *et al.*, 2004), reptiles (Romero *et al*., 2001, Wikelski *et al*., 2003), brush-tailed bettong (*Bettongia penicillata*) (Hing *et al*., 2017), brown bear (*Ursus arctos*) (Cattet *et al*., 2014), and African elephants (*Loxodonta africana*) (Foley *et al*., 2001). Similar to weight, the underlying mechanism of these negative relationships is complex and multifactorial (Cattet, *et al.*, 2014, Cherel, *et al.*, 1988, Foley, *et al.*, 2001, Gray, *et al.*, 1990, Hing, *et al.*, 2017). Therefore, we could not identify whether the effect was due to acute or chronic stressors. Moreover, the sampled koalas were rescued with poor health conditions, which may also affect the relationship between stress and BCS.

#### Limitation and future direction

Due to the nature of this study, there were no healthy koalas admitted. Therefore, one limitation of our study is the lack of healthy koalas for comparison. There was also an unbalanced sample size between different levels, for example, there was only one koala admitted due to trauma. It is unavoidable for research at rescue centers and wildlife hospitals to obtain an unbalanced sample size, and we recommend future hospital-based studies to address this issue by conducting long-term research to increase the overall population size for compensation (Dziura *et al*., 2013). Long-term research could also improve the small sample size in our study, which is reflected in the large standard errors.

Another limitation of this study is that we measured the free cortisol concentration in plasma, which did not include the cortisol bounded to cortisol-binding globulins. In addition, stress induced by capture and transport is also a limitation to this study. The koalas were usually admitted by the public, and inappropriate capturing could result in acute stress. This could be improved by educating the public to contact professionals to capture and transfer wildlife to minimise the stress. The anesthetisation and handling upon arrival for health checks and treatment is another limitation that could induce acute stress in koalas. Ideally, animals should be allowed to habituate to a new environment before handling to minimise stress (Breed *et al*., 2019). However, this is unavoidable in our study because the admitted koalas were under clinical emergencies.

Our study demonstrated that using only one biomarker to assess stress is insufficient to detect and differentiate stressors in settings like rescue centres and wildlife hospitals with no control over the sample population. However, hospital-based studies are critical for koala research due to the high cost of capturing and sampling in the wild (Nyari, *et al.*, 2017). Future studies should investigate the possibility of using more biomarkers to assess stress to reinforce the power of assessment. For example, Beer *et al*. (2023) used an allostatic load index (ALI), which is a quantified index based on multiple biomarkers, to assess cumulative stress in captive giraffes (*Giraffa camelopardalis*).

Allostatic load refers to the dysregulation of the body’s ability to return to homeostasis due to cumulative stressful events (Edes, *et al.*, 2018, Guidi *et al*., 2020). The allostatic load index developed by Beer, *et al.* (2023) included cortisol, DHEAS, cholesterol, non-esterified fatty acids, and fructosamine, which successfully reflected the chronic stress condition of captive giraffes. The use of ALI was rarely done in wildlife (Seeley *et al*., 2022), and we recommended future studies to use our study as a starting point to use multiple biomarkers to establish species-specific ALI to assess chronic stress in wildlife.

In conclusion, our study aimed to identify the use of three biomarkers to measure chronic stress in rescued koalas. No significant difference was found between koalas with different reasons for admission and demographic characteristics. However, the consistency of trends supported by literature found across biomarkers identified Chlamydiosis: reproductive disease, oxalate nephrosis, and conjunctivitis/keratitis as potential chronic stressors. Consistent trends were also identified in other factors, including a higher median in female and adult koalas. Furthermore, negative relationships were found between all biomarkers and age, weight, and BCS, with positive relationships between weight and cortisol and cortisol:DHT as the only exception. This study was limited by the small and unbalanced sample size and a lack of a healthy population due to the nature of hospital-based studies. Stress induced by capture, transport, and handling was another limitation of our study. Long-term studies should be conducted in the future to compensate for the sample population issue. This finding has important implications for the potential use of cortisol:DHEAS and cortisol:DHT to detect stress. Moreover, our study identified the insufficiency of using a single biomarker to assess chronic stress and established a foundational framework that future studies could use to develop koala-specific ALI to understand chronic stress in rescued koalas.

## Data availability

The data and R code underlying this article will be made available upon request via email to the corresponding author.

## Acknowledgements

We acknowledge and pay our respect to the Traditional Owners and Custodians of the lands where we work that belong to the Yuggera peoples. We also acknowledge the veterinarians at RSPCA Queensland for the sample and data collection and the stress lab for all the assistance and support throughout this study.

## Conflicts of interest

The authors have no conflicts of interest to declare.

## Notes

### Competing Interest Statement

The authors have declared no competing interest.

